# SBE1 drives the circumferential growth of the starch sheath around the *Chlamydomonas reinhardtii* pyrenoid

**DOI:** 10.64898/2026.07.31.742166

**Authors:** Micah I. Burton, Haoyu Wu, Lianyong Wang, Jessica H. Hennacy, Martin C. Jonikas

## Abstract

Pyrenoids are CO2-fixing organelles responsible for approximately one-third of global CO2 fixation. The pyrenoids of many algae are surrounded by a starch sheath proposed to perform the critical function of slowing leakage of concentrated CO2 out of the pyrenoid. How the cell shapes starch granules into curved starch plates that encase the pyrenoid to enable efficient CO2 fixation is currently unknown. Here, we elucidate how starch transitions from granules into a fully formed pyrenoid starch sheath using confocal microscopy of the model green alga *Chlamydomonas reinhardtii*. We observe that after initiation, starch granules grow circumferentially along the surface of the pyrenoid matrix to encapsulate it. We show that the starch branching enzyme SBE1 localizes to the pyrenoid and is essential for this circumferential starch granule growth. Our data suggest that SBE1 promotes the circumferential growth of pyrenoid-associated starch granules by branching starch at the matrix-granule-stroma interface. Our findings advance the understanding of pyrenoid starch sheath assembly and, more broadly, starch-shaping mechanisms.

**Significance Statement:** The shape of the starch granules that surround the pyrenoid is critical for efficient carbon fixation, but how these granules are shaped into curved plates is currently not understood. We find that starch granules spread along the pyrenoid surface rather than growing equally in all directions, and that this bias requires SBE1, a branching enzyme localized to the surface of the spherical pyrenoid matrix condensate. The findings support a model in which localizing a metabolic enzyme directs polymer growth to create curvature. This mechanism connects branching enzyme placement to starch granule geometry and contributes to the understanding of how cells convert local biosynthetic activity into large-scale organelle architecture.

## Introduction

Pyrenoids, cellular organelles that enhance carbon fixation, mediate approximately one-third of global CO2 fixation (1). Pyrenoids are found in eukaryotic algae, as well as non-vascular land plants belonging to the hornworts lineage (2, 3). Recent work characterizing the pyrenoid has sought not only to understand its basic biology (4–10) but also to enable its engineering into major crop plants to increase yield for societal benefit (6, 11, 12).

The most comprehensive molecular understanding to date of pyrenoid assembly and function has been obtained in the model green alga *Chlamydomonas reinhardtii* (hereafter, Chlamydomonas) (13), owing to its genetic tractability and molecular toolkit (14, 15). In Chlamydomonas, the pyrenoid is composed of three sub-compartments: a phase-separated condensate (5, 8, 9, 16) primarily composed of the CO2-fixing enzyme Rubisco (17), membrane tubules traversing the organelle (6, 18, 19) thought to deliver CO2 to Rubisco for fixation (20, 21) following uptake (22, 23), and a sheath composed of starch that surrounds the spherical matrix (18). The starch sheath has been proposed to serve both as a barrier to slow CO2 escape from the matrix (24) and to act as a scaffold for proteins involved in pyrenoid function (25).

The starch sheath is built from starch, a glucose polymer found throughout the *Chloroplastida*, a lineage that includes green algae and all land plants (26). This ubiquitous polymer is primarily used for energy storage and consists of two classes of α-glucans: amylopectin, which typically makes up the majority of a starch polymer (>70%) and consists of linear α-1,4 glycosidic linkages that contain ∼5% α-1,6 branch points, and amylose, which constitutes the remainder, and has the same composition as amylopectin but branches very rarely (27).

While the chloroplast stroma of leaves (28) and green algae (29) usually contain lenticular starch granules, the granules that make up the pyrenoid starch sheath adopt a dramatically different geometry. To form the sheath, Chlamydomonas shapes starch granules into curved plates that closely encapsulate the pyrenoid matrix (18, 24, 30). Given that cells can synthesize either lenticular starch granules in the stroma or shell-like granules around the pyrenoid matrix, there are likely specific molecular mechanisms that determine starch granule morphology. However, these mechanisms remain largely unknown (31). Thus, understanding how the starch sheath forms around the pyrenoid matrix can provide key insights not only into the establishment of pyrenoid function but also into general starch shaping mechanisms.

In this work, we contribute to the understanding of starch sheath formation by observing the real-time assembly of the starch sheath upon transition from high to low CO2 and after cell division, and by characterizing a mutant with impaired starch sheath formation. Our data show that Starch Branching Enzyme 1 (SBE1; Cre06.g289850) contributes to the shaping of pyrenoid starch granules into a starch sheath by promoting the circumferential growth of starch granules around the pyrenoid matrix. Together, our findings help to elucidate how the pyrenoid starch sheath is shaped and suggest a novel mechanism by which a starch branching enzyme can influence starch shaping.

## Results

### Live imaging of Chlamydomonas by confocal microscopy highlights starch granule localization and dynamics during starch sheath formation

To characterize how starch is assembled into the plates that surround the pyrenoid of Chlamydomonas cells, we sought to observe starch granules in individual cells over time as the starch sheath formed. Previous work found that the Chlamydomonas starch sheath formed as cells acclimated from high levels of CO2 to low levels of CO2 during the induction of the CO2-concentrating mechanism (24, 30). However, due to the technical limitations of observing fixed cells, these prior studies could not track starch granules within a single cell over time. To improve the understanding of how starch granules are formed into the starch sheath, we visualized starch granules in otherwise wild-type Chlamydomonas cells using the previously-identified starch marker granule-bound starch synthase IA (GBSS1A, Cre17.g721500; STA2) tagged with the fluorescent protein Venus (4). We imaged live cells over a time course of 480 minutes using confocal microscopy during starch sheath assembly upon transition from photoautotrophic growth at high (3% v/v) levels of CO2 to low (0.04% v/v) levels of CO2 (Fig. 1*A*).

**Figure 1.**
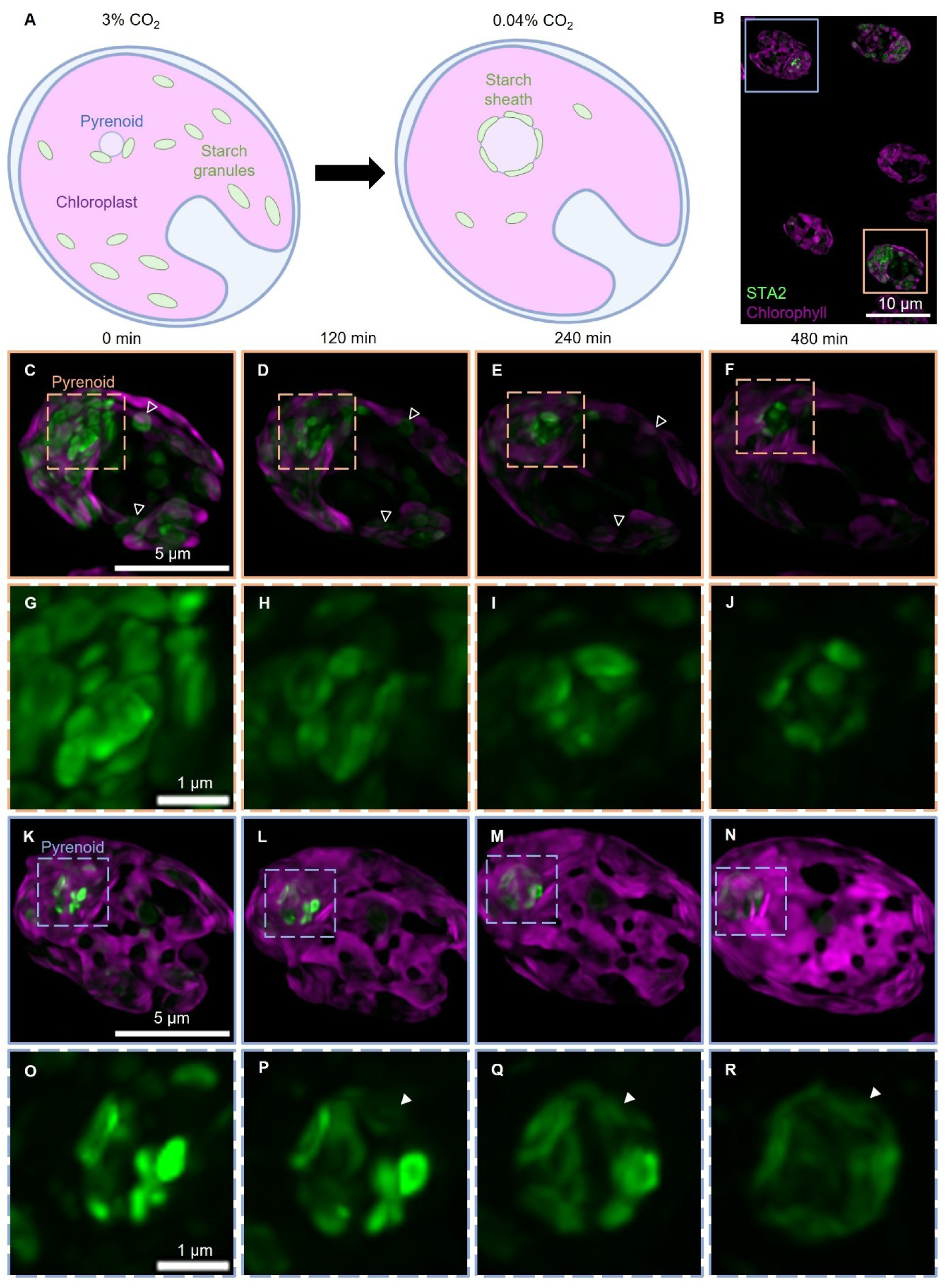
Upon transfer to low CO2, the pyrenoid starch sheath is formed from proximal starch granules. (A) Cartoon showing starch distribution in a cell acclimated to high levels of CO2 (3% v/v) or low (0.04% v/v) levels of CO2. (B-R) Cells expressing the starch marker STA2-Venus were grown in TP minimal medium at high CO2 and transferred to a microscopy slide covered with a TP-agarose pad at low levels of CO2. Cells were supplied with continuous illumination. Images were acquired every 15 minutes. (B) Confocal Max-Z projection of cells immediately after transfer to air levels of CO2. (C-R) Confocal Max-Z projections of cells acclimating to air levels of CO2. (C-F) Time course of a representative high-to-low CO2-acclimating cell with large amounts of starch distributed throughout the cell (the cell in the orange box in panel B). Empty arrowheads denote stromal STA2 signal that is decreasing in intensity. The dashed-boxed region represents the canonical pyrenoid location. (G-J) Zoom-in of the dashed-boxed region in (C-F) showing only STA2-Venus signal. (K-N) Time course of a representative high-to-low CO2-acclimating cell with starch distributed primarily around the pyrenoid. The boxed region represents the canonical pyrenoid location. (O-R) Zoom-in of the boxed region in (K-N) showing only STA2-Venus signal. Arrowheads indicate the location of a growing starch plate.

Initially, high CO2-acclimated cells contained varying amounts of starch granules with no clear starch sheath, as evidenced by the lenticular-shaped STA2-Venus fluorescence present throughout the chloroplast of cells (Fig. 1 *B, C* and *K*). In cells that contained large numbers of starch granules, starch appeared to be distributed throughout the chloroplast stroma (Fig. 1*C*). In contrast, cells that contained fewer granules tended to have starch localized primarily to a region within the cell consistent with the canonical location of the pyrenoid (Fig. 1*K*). As cells acclimated to low levels of CO2, we observed that the STA2-Venus signal was retained in the canonical pyrenoid region while the STA2-Venus signal decreased in size and brightness in the stroma (Fig. 1 *D-F*, *H-J*, *L-N*, *P-R* and Movie S1). Our observations are in agreement with previous studies that observed localized degradation of starch in the stroma during the high-to-low CO2 transition (30).

We observed a slight decrease in the brightness of STA2-Venus signal in the pyrenoid region at time points exceeding 240 minutes (Fig. 1 *F, J, N, R* and Movie S1); however, starch granules still grew as judged by an increase in the extent of STA2-Venus signal (Fig. 1 *P-R*, arrowheads). This decrease in signal could be the result of the photobleaching of older starch plates which are no longer growing quickly, as STA2 is incorporated into starch granules as they grow (32). We conclude that our STA2-Venus time course recapitulates the previously-known starch degradation in the stroma and starch accumulation around the pyrenoid during the acclimation from high to low levels of CO2.

### Starch plates are inherited during cell division and reused in the starch sheaths of daughter cells

During cell division, the pyrenoid matrix is distributed to daughter cells by either fission, dissolution, or inheritance of the matrix to a single daughter (8, 33). What happens to the starch sheath in this context is poorly understood. In our high-to-low CO2 time course experiments, we fortuitously observed several dividing cells that allowed us to observe how the starch sheath is distributed to daughter cells during cell division. We observed that the starch sheath plates were disrupted before the cleavage furrow bisected the chloroplast, as judged from autofluorescence (Fig. 2 *B-D*, white arrowheads). This detachment could potentially be promoted by dissolution of the matrix, which occurs before cell division (8, 34), or from the physical forces induced by the approaching cleavage furrow disrupting the pyrenoid as observed in a previous study (33). As the cleavage furrow encroached on the pyrenoids, we observed distribution of the starch plates to either side of the furrow, depending on the position they were in initially (Fig. 2 *E-G*). Once complete cleavage occurred, we observed that the starch plates inherited by daughter cells were repositioned, presumably surrounding the pyrenoid matrix in each cell (Fig. 2 *I-N* and Movie S2). We conclude that the starch plates that compose the sheath are inherited during cell division by daughter cells depending on each granule’s location relative to the cleavage furrow and are reused in the pyrenoids of the daughter cells.

**Figure 2.**
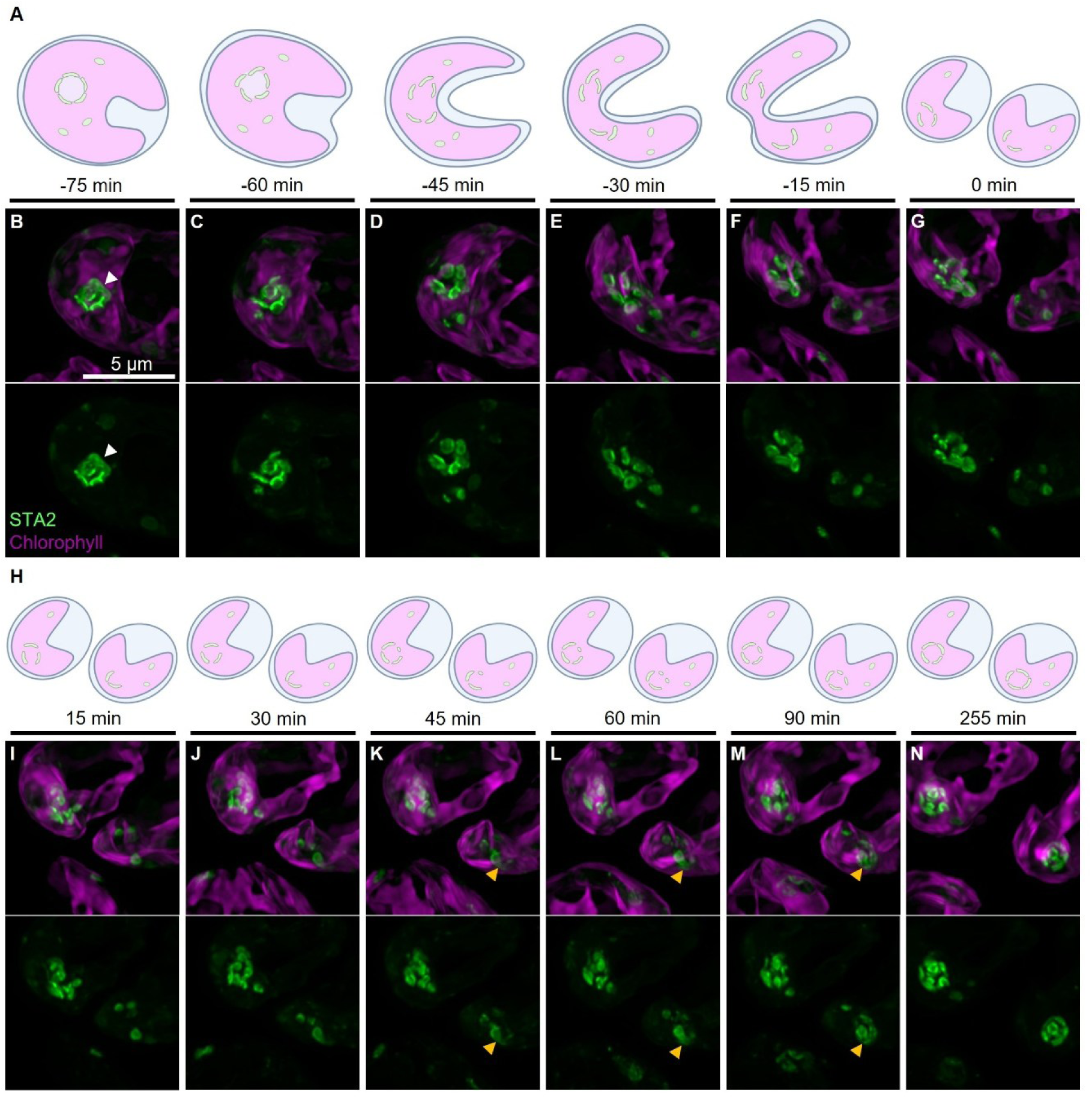
During cell division, cells inherit starch plates and grow new granules between inherited plates. (A-N) Representative time course of cell division of a cell expressing STA2-Venus grown in TP minimal medium at high (3% v/v) CO2 then transferred to a microscopy slide covered in a TP-agarose pad at air (0.04% v/v) levels of CO2 for 5 hours prior to the first acquisition. Cells were supplied with constant illumination. (A) Schematic of cleavage furrow migration based on chlorophyll autofluorescence for panels (B-G). (B-G) Confocal Max-Z projections of the pyrenoid leading up to cell division. White arrowheads denote the location of the starch sheath. t = 0 min is the first observed instance of complete separation of chlorophyll signal. (Bottom) Venus-only channel of panels (B-G). (H) Schematic of starch sheath development in the chloroplast for panels (I-N). (I-N) Confocal Max-Z projections of the pyrenoid region of two daughter cells after complete chloroplast division. Yellow arrowheads denote a daughter pyrenoid where both starch granules from the mother cell are reused in the formation of the starch sheath and starch granules appear to be formed *de novo* in locations around the daughter pyrenoid that lack starch. (Bottom) Venus-only channel of panels (I-N).

In our observations, one daughter cell often inherited the majority of the starch plates and the other inherited few or no starch plates (Fig. 2*G*). When this occurred, even cells that inherited too few plates to fully surround the pyrenoid still assembled a nearly complete starch sheath within 90-180 minutes (Fig. 2 *I-N* and Movies S2 and S3). The gaps between the inherited granules appeared to be filled via the growth of new granules or the incorporation of nearby nascent starch granules (Fig. 2 *K-M*, yellow arrowheads). We conclude that daughter cells can rapidly assemble a new starch sheath by using the starch plates inherited from mother cells or by synthesizing new starch granules.

### Pyrenoid-associated starch granules grow circumferentially to fully surround the pyrenoid

To better visualize the growth of starch granules around the pyrenoid, we sought to reduce the amount of starch present around the pyrenoid. Starch accumulation is thought to be controlled by the circadian clock in Chlamydomonas cells (32). When cells are grown autotrophically under a 12-hour-light/12-hour-dark diurnal cycle, starch accumulates during the light period and is degraded during the dark period (35, 36). After acclimating cells to such a cycle, we shortened the final light period to 8 h and extended the subsequent dark period to 16 h. We reasoned that this treatment would reduce starch levels at the start of imaging, as previously shown in *Arabidopsis thaliana* (37).

We imaged diurnally entrained cells using confocal microscopy, beginning right after the dark to light transition at the end of the extended 16-hour night. We observed that initially, cells had incomplete starch sheaths, as visualized by STA2-Venus signal (Fig. 3*A*). As cells grew in the presence of light, the initial starch granules appeared to grow circumferentially from their origin points, reaching near full coverage of the pyrenoid matrix in approximately 120 minutes (Fig. 3 *A-D*, white arrowheads and Movie S4) and complete coverage by 240 minutes (Fig. 3*E* and Movie S4). This observation agrees with the previously reported rapid accumulation of starch in diurnally entrained cells during the first 2 hours of the light period (36).

**Figure 3.**
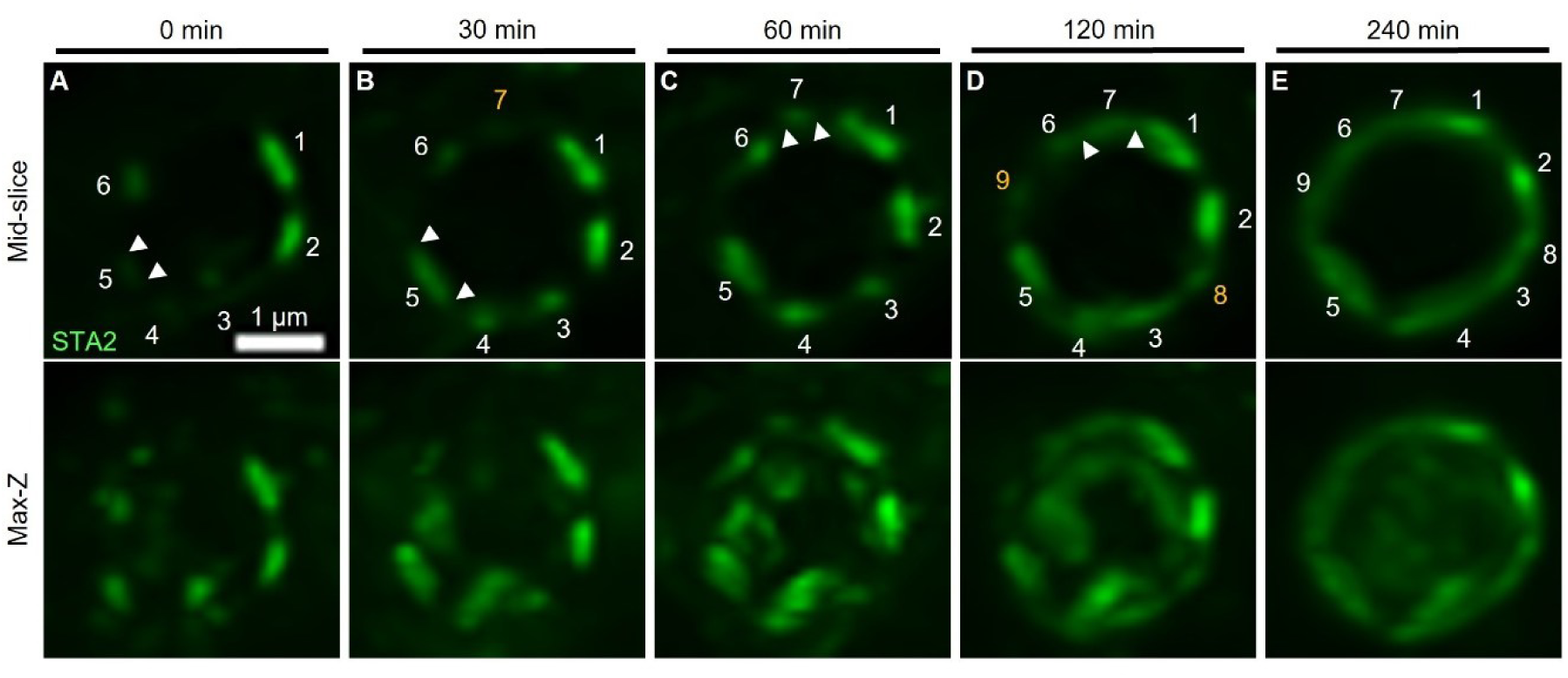
Starch granules grow circumferentially along the surface of the pyrenoid. (A-E) Confocal Max-Z projection of 2 Z-slices (“Mid-slice”) which pass through the center of the pyrenoid, or full Max-Z projection (“Max-Z”) of a representative wild-type cell expressing STA2-Venus. Cells were grown in TP minimal medium at air-levels of CO2 under a 12h-light/ 12h-dark cycle for at least 7 days and then subjected to one 8h-light/16h-dark cycle prior to imaging (0 min). Cells were imaged on TP-agarose pads and supplied with constant illumination. Images were acquired every 15 minutes. White numbers indicate individual starch plates. Yellow numbers indicate nascent starch plates. Arrowheads denote the edges of starch granules.

Cells also appeared to synthesize new starch granules with progression of time in the light. In cells where the initial starch granule distribution around the pyrenoid was sparse, we observed what appeared to be the *de novo* growth of starch granules between growing starch plates (Fig. 3 *B-D*, yellow numbers). One caveat to this finding is that these granules may initially have been too small to detect with our equipment or moved between acquisitions; however, we still observed a preference for the circumferential growth of these granules (Fig. 3 *B-E* and Movie S4). We conclude that the starch sheath originates from circumferentially growing starch granules that are either retained from a previous sheath or are initiated on the pyrenoid matrix surface.

### SBE1 is required for the circumferential growth of the starch sheath around the pyrenoid matrix

To identify putative proteins involved in promoting the circumferential growth of the starch sheath around the pyrenoid, we looked at starch-related proteins that were previously found to interact with known pyrenoid proteins (4). One of these proteins, SBE1, was previously found to interact with an isoform of the small subunit of Rubisco (RBCS2; Cre02.g120150) (4). SBE1 is predicted to contain three functional domains common to all starch branching enzymes (38) (Fig. 4*A*). These domains include a carbohydrate-binding module 48 domain at the N-terminus, a centrally located glycosyl hydrolase 13 family catalytic domain, and an α-amylase family associated all-β domain at the C-terminus (38). In addition, SBE1 is also predicted to contain four Rubisco-binding motifs that may mediate interaction with RBCS2 (5) (Fig. 4*A*). Previous work characterized an *sbe1* mutant and found it had a starch catabolism defect under nitrogen starvation conditions (39). A more recent study reported no significant changes in starch composition in *sbe1* mutant cells under the same conditions and heterotrophic growth (38). Neither study characterized an *sbe1* mutant under autotrophic growth. This led us to reason that the enzyme may play a non-essential role in overall starch metabolism and instead operate specifically in the context of starch sheath development.

**Figure 4.**
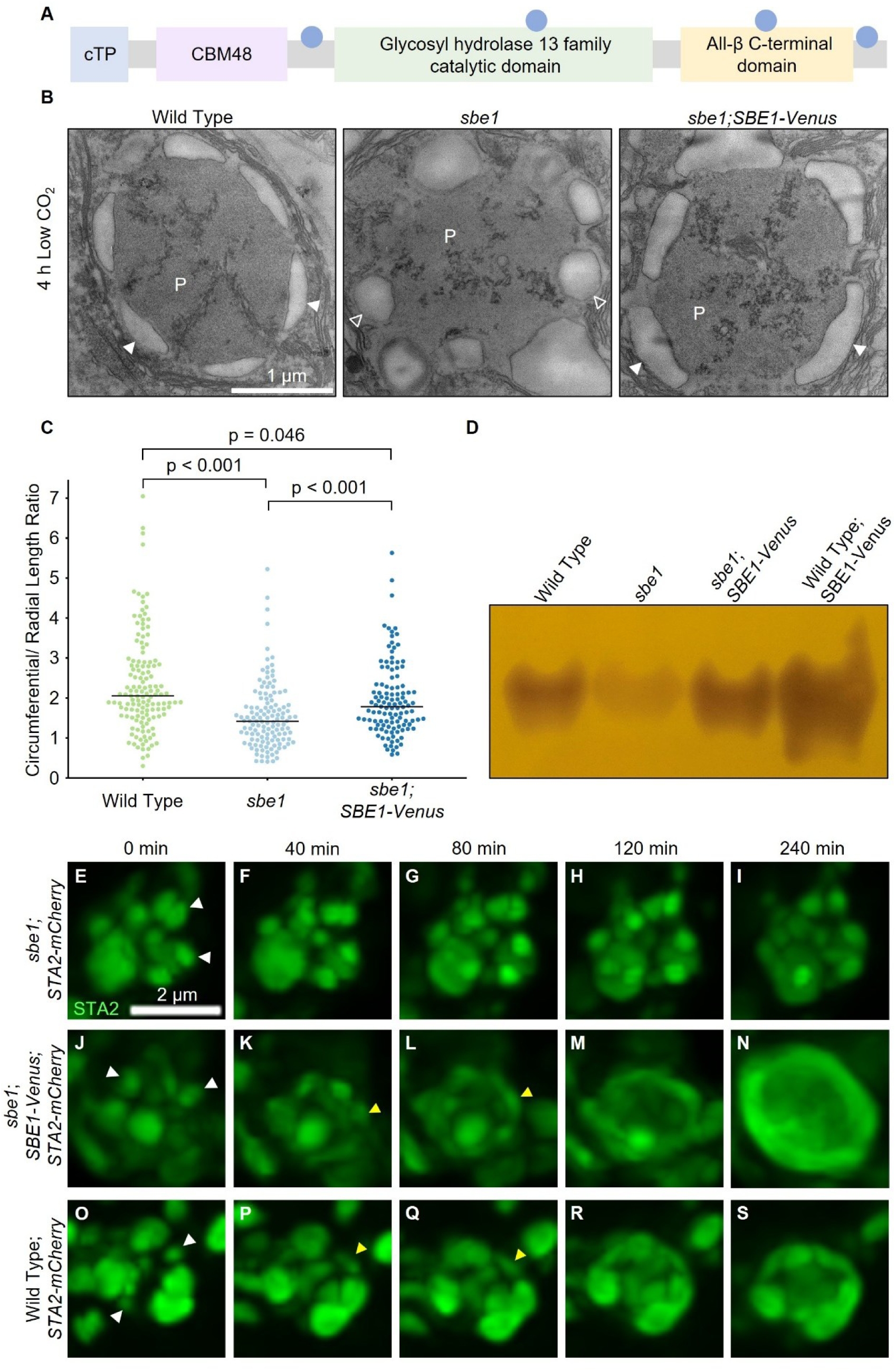
SBE1 is required for circumferential growth of starch plates during acclimation from high (3% v/v) to low (0.04% v/v) CO2 levels. (A) Cartoon of the domain structure of SBE1. cTP - chloroplast transit peptide. CBM48 - carbohydrate-binding module 48. Circles indicate the relative locations of putative Rubisco-binding motifs (5). (B) Representative transmission electron microscopy images of the pyrenoids of wild-type, *sbe1* mutant, and *sbe1;SBE1-Venus* cells harvested 4 hours after a shift from high to low levels of CO2 in TP minimal media. Solid arrowheads denote starch sheath plates. Empty arrowheads denote abnormally-shaped starch sheath granules. P indicates the pyrenoid. (C) Quantification of the ratio between the circumferential (the arc length a given granule covers when in contact with the pyrenoid matrix) and radial (the perpendicular length of the granule from the midpoint of the circumferential length) lengths of pyrenoid matrix associated starch granules in wild-type (n = 137), *sbe1* mutant (n = 128), and *sbe1;SBE1-Venus* (n = 116) cells grown in TP minimal medium at 4 h low (0.04% v/v) CO2. *P-*values were determined by using a Kruskal-Wallis test (χ^2^ = 41.87, d*f* = 2, *P* = 8.093 × 10^-10^) followed by Dunn’s multiple comparisons test with Bonferroni correction. Horizonal bars indicate the median of the distribution. (D) Zymogram using a native gel containing no polysaccharide showing starch branching activity from lysate harvested from wild-type, *sbe1* mutant, *sbe1;SBE1-Venus*, and wild-type;*SBE1-Venus* cells grown in high CO2 and shifted to low CO2 for 21 hours in TP minimal media. 50 µg of protein was loaded in each labeled well. The gel was incubated for 16 hours at 4°C in assay buffer containing phosphorylase A with gentle shaking prior to staining with Lugol’s solution (1% KI, 0.1% I2). (E-S) Confocal microscopy max-Z projection images of the pyrenoids of representative *sbe1* mutant (E-I), *sbe1;SBE1-Venus* rescue (J-N), and wild-type (O-S) cells expressing STA2-mCherry grown at low CO2 levels under continuous illumination. Cells were grown at high CO2 in liquid TP minimal medium until time point zero. Images were acquired every 20 minutes. White arrowheads denote starch granules that were initially present around the pyrenoid matrix. Yellow arrowheads denote circumferentially growing starch granules.

To test if SBE1 is necessary for assembly of the starch sheath, we obtained a mutant in the SBE1 gene from the CLiP mutant library (LMJ.RY0402.151042) (14) and verified the presence of the insertional cassette using PCR (*SI Appendix*, Fig. S1). We then cultured wild-type and *sbe1* mutant cells under autotrophic conditions in high levels of CO2 to degrade the starch sheath before transferring them to low levels of CO2 to induce the formation of the starch sheath. After 4 hours of growth under low-CO2 conditions to capture the final stages of starch sheath development (24, 30), we fixed the cells and prepared them for TEM imaging. We observed that in *sbe1* mutant cells, the starch granules that normally formed a shell encasing the pyrenoid matrix lacked the characteristic starch sheath plate-like morphology (Fig. 4*B* and *SI Appendix*, Figs. S2 and S3). Measurements of the circumferential-to-radial length ratio of each pyrenoid matrix-associated starch granule revealed that the starch granules in *sbe1* mutant cells were significantly more spherical, with a median circumferential-to-radial length ratio of 1.42, as compared to wild-type cells, which were more elongated circumferentially and had a median ratio of 2.06 (*P* < 0.001, Dunn’s multiple comparisons test with Bonferroni correction; Fig. 4*C*). This suggested to us that SBE1 potentially plays a role in promoting the circumferential growth of starch granules around the pyrenoid matrix.

While we observed a clear phenotypic difference in starch sheath morphology between wild-type cells and *sbe1* mutant cells, CLiP mutants have the potential to harbor off-target insertion sites, which could be responsible for the observed phenotype (15). Therefore, we genetically rescued the *sbe1* CLIP mutant using a construct that produced SBE1 fused to the fluorescent protein Venus followed by a 3xFLAG tag under the native promoter of the *SBE1* gene (hereafter, SBE1-Venus). We confirmed that cells expressing SBE1-Venus produced catalytically active protein using native zymography. We observed that while the lysates from *sbe1* mutant cells showed a faint band of branching activity, wild-type and *sbe1;SBE1-Venus* cells showed increased branching activity as determined by increase staining (Fig. 4*D* and *SI Appendix*; Fig. S5*A*). Additionally, wild-type cells that expressed SBE1-Venus showed greater staining than either wild-type or *sbe1;SBE1-Venus* (Fig. 4*D* and *SI Appendix*; Fig. S5*A*). We conclude that the *sbe1* mutant lacks a starch branching activity that is rescued by exogenous expression of SBE1-Venus.

Previous work showed additional bands of activity when performing native zymography on lysates harvested from cells grown in heterotrophic conditions (TAP media), which we failed to detect in our zymograms carried out on cells grown under autotrophic conditions (TP minimal media) (38). We observed that heterotrophically-grown wild-type cells displayed additional branching activity that was not detected when the cells were grown under autotrophic conditions (*SI Appendix*; Fig. S5*B*). This suggests that Chlamydomonas cells may produce either varying concentrations of branching enzymes, branching activity is modulated, or that branching enzymes are regulated into different complexes depending on growth conditions.

To identify if expression of SBE1-Venus could rescue the phenotype of the *sbe1* mutant, we performed TEM on *sbe1;SBE1-Venus* cells grown autotrophically under high CO2 before shifting to growth at low CO2 for 4 hours prior to harvest and measured the circumferential-to-radial length ratio of the pyrenoid matrix-associated starch granules. We found that instead of the spherical pyrenoid matrix-associated starch granules that we observed in the *sbe1* mutant (median ratio 1.42), the *sbe1;SBE1-Venus* cells grew more elongated starch plates and had a median ratio of 1.78 (*P* < 0.001, Dunn’s multiple comparisons test with Bonferroni correction; Fig. 4 *B* and *C* and *SI Appendix*, Figs. S3 and S4). Our *sbe1;SBE1-Venus* cells did not fully recover the circumferential-to-radial length ratio of wild-type cells (median ratio of 1.78 in *sbe1;SBE1-Venus* cells vs. 2.06 in wild-type cells; *P* = 0.046, Dunn’s multiple comparisons test with Bonferroni correction; Fig. 4 *B* and *C* and *SI Appendix*, Figs. S2 and S4), potentially due to lower *in vivo* activity of SBE1-Venus.

We next sought to observe how the abnormal starch granules in *sbe1* mutant cells were formed and how the expression of our SBE1-Venus construct rescued their morphology. We transformed a construct that contained STA2 tagged at the C-terminus with mCherry into our *sbe1, sbe1;SBE1-Venus,* and wild-type lines and observed the formation of the starch sheath as the cells acclimated from high to low levels of CO2, the same conditions as used in preceding experiments (Fig. 1 *B-R* and 4*B*). When acclimated to high CO2 levels, *sbe1* mutant, *sbe1;SBE1-Venus*, and wild-type cells each contained spherical starch granules that were distributed around the pyrenoid matrix (Fig. 4*E, J* and *O,* white arrowheads). As the cells acclimated to low levels of CO2, the morphology of the starch granules in the *sbe1* mutant cells remained relatively unchanged throughout the entire time course (Fig. 4*E*-*I* and Movie S5). In contrast, the initial starch granules around the pyrenoids in *sbe1;SBE1-Venus* and wild-type cells grew circumferentially (Fig. 4*K, L*, *P,* and *Q,* yellow arrowheads and Movies S6 and S7). Eventually, these growing starch granules formed complete starch sheaths (Fig. 4*M, N, R,* and *S* and Movie S6 and S7). Given that SBE1-Venus restores the circumferential synthesis of starch around the pyrenoid matrix in *sbe1* mutant cells, we conclude that SBE1 is necessary for the circumferential growth of the starch granules around the pyrenoid matrix during the high-to-low CO2 transition.

### The starch-branching enzyme SBE1 localizes to the pyrenoid matrix-starch interface

To determine how SBE1 contributes to the circumferential growth of starch around the pyrenoid matrix, we sought to determine where the protein localized in the cell. Therefore, we imaged *SBE1-Venus* cells using confocal microscopy. We found that SBE1-Venus signal was enriched as a ring at the canonical pyrenoid location within the cell, reminiscent of the starch sheath (Fig. 5*A*). To test if the localization pattern of SBE1 corresponded with the starch sheath, we co-expressed *SBE1-Venus* along with *STA2-mCherry* in *sbe1* mutant cells and found that both proteins localized in the same ring pattern (Fig. 5*B*).

**Figure 5.**
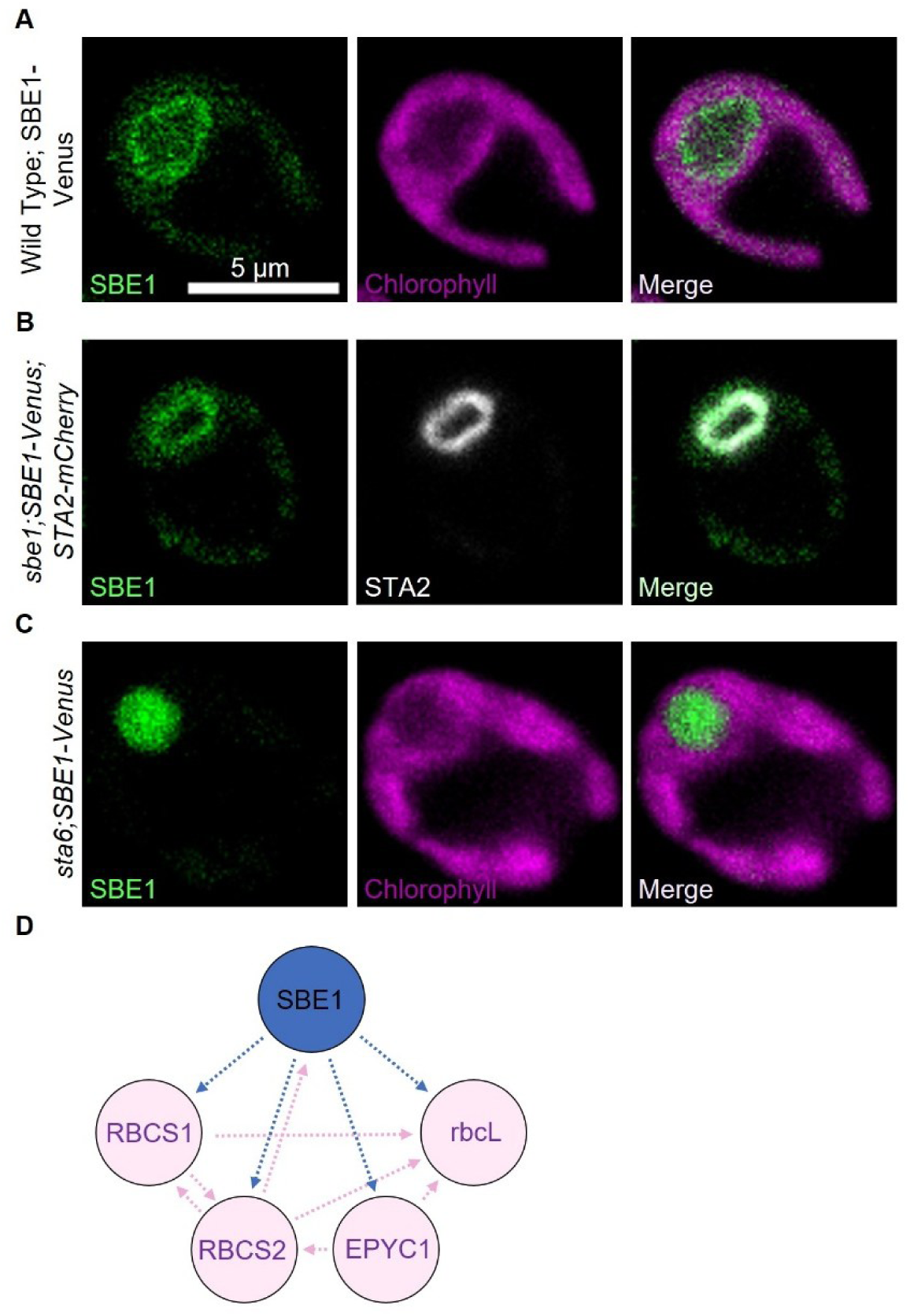
The pyrenoid matrix-localized starch branching enzyme SBE1 is targeted to the pyrenoid matrix periphery in the presence of starch. (A-C) Confocal single Z-slices of representative Chlamydomonas cells expressing SBE1-Venus. Cells were grown in high (3% v/v) CO2 and transferred to low (0.04% v/v) CO2 overnight before imaging. (A) Wild-type cells. (B) *sbe1* mutant cells additionally expressing STA2-mCherry. (C) *sta6* mutant cells. (D) Diagram depicting enriched pyrenoid matrix proteins that were detected by mass spectrometry following co-immunoprecipitation using SBE1-Venus-3xFLAG as bait. Arrows extend from bait proteins to prey proteins. Magenta arrows are protein interactions that have been identified previously (4). Blue arrows indicate novel protein interactions from this study.

In some cells, we observed that SBE1 did not localize to the periphery of the pyrenoid but instead to the pyrenoid matrix. We hypothesized that this could be the result of these cells lacking a starch sheath. To test if this alternative localization was indeed due to a lack of the starch sheath, we transformed the Chlamydomonas *sta6* starchless mutant CC-5374, deficient in the catalytic small subunit of ADP-glucose pyrophosphorylase (40), with SBE1-Venus and imaged the cells using confocal microscopy. We found that in the starchless mutant background, SBE1-Venus no longer localized to the periphery of the pyrenoid and instead localized solely to the canonical location of the pyrenoid matrix (Fig. 5*C*). Together, our data suggest that SBE1 localizes to the pyrenoid matrix-starch interface in the presence of starch, and to the pyrenoid matrix in the absence of starch.

### SBE1 physically interacts with pyrenoid matrix proteins

It has been reported that starch branching enzymes can form complexes with other proteins involved in starch synthesis to carry out starch anabolism (41). Therefore, we sought to determine if SBE1 forms complexes with other enzymes involved in starch assembly. To find interaction partners, we carried out immunoprecipitation followed by mass spectrometry analysis using wild-type cells expressing SBE1-Venus-3xFLAG. Our pull-down reproduced the previously known interaction between RBCS2 and SBE1 (4) using SBE1 as bait instead of RBCS2 as in the previous study and revealed novel interactions with key components of the pyrenoid matrix including RBCS1 (Cre02.g120100), rcbL, and EPYC1 (Cre10.g436550) as determined by their enrichment above the Venus control (Fig. 5*D* and Dataset S1). Considering that SBE1 contains four potential Rubisco-binding motifs (5) (Fig. 4*A*), SBE1 could be pulling down the Rubisco holoenzyme and EPYC1 thorough its interaction with the Rubisco small subunits rather than directly interacting with each pyrenoid matrix component. We did not find interactions with other proteins involved in starch synthesis or degradation that were consistently reproduced or enriched above the Venus control, suggesting that SBE1 either acts alone, the Venus-tag interferes with interactions with other proteins, or SBE1 forms transient interactions that could not be detected with our approach. Considering SBE1’s predicted Rubisco-binding motifs (5) (Fig. 4*A*), our finding that SBE1 localizes solely to the pyrenoid matrix in the absence of starch (Fig. 5*C*), and the interaction we identified between SBE1 and pyrenoid matrix components (Fig. *5D* and Dataset S1), our data suggest that SBE1 is localized to the pyrenoid through its interaction with Rubisco.

## Discussion

Previous efforts to study starch sheath formation used either TEM (24) or light microscopy (30) to observe cellular populations at discrete timepoints during the acclimation of cells from high to low CO2 conditions. These studies were limited in that they did not follow individual granules and as such, the proposed mechanisms for starch sheath formation were largely speculative in nature. To overcome these limitations, here, we used confocal microscopy to visualize the growth of individual starch granules in live cells. By observing starch sheath development during the acclimation of cells from high to low levels of CO2 (Fig. 1 and Fig. 4 *O-S*) and during formation of the starch sheath in cells already acclimated to low levels of CO2 after an extended night (Fig. 3), we found that the initial starch sheath granules were either already present even at high levels of CO2 (Fig. 1 *K* and *O* and Fig. 4*O*), or were apparently formed *de novo* on the surface of the pyrenoid matrix (Fig. 3). In both cases, we observed that the initial starch granules grew in a circumferential manner to surround the pyrenoid matrix (Fig. 3 and Fig. 4 *O-S*).

A prevailing hypothesis suggests that the starch sheath forms due to the sequestration of Rubisco to a single compartment within the chloroplast, and thus starch deposition along the pyrenoid matrix occurs as the result of a gradient of 3-PGA or other metabolites produced from localized carbon fixation (26, 29, 30, 42). Challenging this view, recent work characterizing the starch-binding proteins StArch Granules Abnormal 1 (SAGA1, Cre11.g467712) and SAGA2 (Cre09.g394621) in Chlamydomonas (10) and another study demonstrating the recruitment of starch to a Rubisco-EPYC1 matrix in an *A. thaliana* line engineered to express SAGA1 and SAGA2 (11) have shown that these proteins are both necessary and sufficient to induce the formation of starch granules on the surface of the pyrenoid matrix. Here, we show that the pyrenoid-localized starch-branching enzyme SBE1 is required for the circumferential growth of starch sheath granules (Fig. 4*A*, Fig. 5 *A-C*), as *sbe1* mutant cells were unable to form starch plates from initial starch granules (Fig. 4*B* and *C* and *E-I*). We were able to rescue the *sbe1* mutant phenotype and restore the circumferential growth of starch sheath granules with the expression of an SBE1-Venus construct (Fig. 4*B*, *C* and *J-N*). Our findings demonstrate that starch granule localization to the surface of the pyrenoid matrix and the shaping of the pyrenoid starch sheath granules into their canonical shape can be uncoupled in Chlamydomonas.

We propose a model in which SBE1 promotes the circumferential growth of starch around the pyrenoid (Fig. 6). Initially, in the absence of a starch sheath, SBE1 localizes to the pyrenoid matrix, likely by binding to Rubisco (Fig. 5 *C* and *D* and Fig. 6*A* and Dataset S1). Once starch synthesis increases at the matrix surface during the high-to-low CO2 transition, possibly by factors promoting starch synthesis and recruitment such as SAGA1 and SAGA2 (10, 11), SBE1 is recruited to the starch-matrix interface possibly due to the affinity of the enzyme for both starch and the matrix (Fig. 5 *B* and *D* and Fig. 6 *A-B*). SBE1 then acts to branch starch along the starch-matrix interface providing stromal starch synthases newly branched starch ends to extend linearly at the intersection of the matrix, starch, and stroma (Fig. 6*B*, blue branches). Each enzymatic activity fuels the other, functionally mirroring primer-free starch branching zymography (43–45) (Fig. 4*D* and *SI Appendix*; Fig. S5). SBE1 behaves like a type 1 starch branching enzyme (38), which prefers to branch long, linear chains of starch (38, 46). Therefore, SBE1 activity is likely favored at the starch-matrix-stromal edges of pyrenoid associated starch granules, where stromal starch synthases can access and extend starch ends, resulting in the circumferential growth preference of starch plates in wild-type cells (Fig. 6 *B* and *C*, blue arrows). In the absence of SBE1, any starch granules that form on the surface of the pyrenoid would only be branched by stromal starch branching enzymes, resulting in similar rates of circumferential and radial growth, and thus explaining the more rounded cross-sections, of the pyrenoid-associated granules present in *sbe1* mutant cells (Fig. 4*B*, *C* and *E-I* and Fig. 6*D*-*F*).

**Figure 6.**
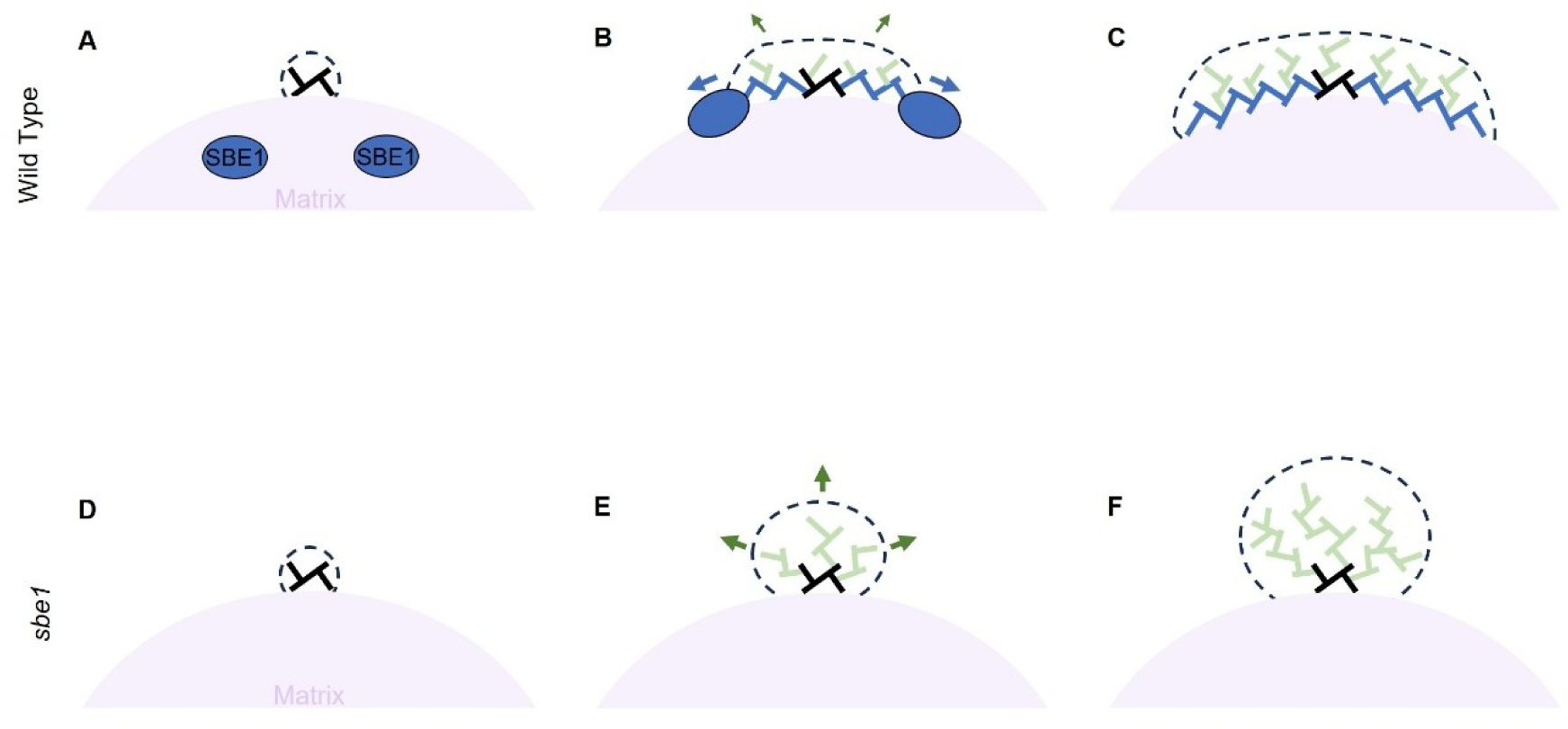
Model: SBE1 promotes the circumferential growth of starch granules at the surface of the pyrenoid. (A-F) Model of starch plate growth in either wild-type cells (A-C) or *sbe1* mutant cells (D-F). Black branches represent initial starch formed on the surface of the matrix. Blue branches represent starch produced through SBE1 activity on the growing starch plate. Green branches represent stromal starch branching activity. Arrows indicate the direction of starch growth promoted by each branching activity (green for stromal and blue for SBE1). Dashed lines outline the starch granule.

Our discovery of the role of SBE1 in pyrenoid starch sheath shaping contributes to efforts to engineer a pyrenoid into land plants to increase crop yields and climate resilience. A condensate composed of EPYC1 and Rubisco (proto-pyrenoid) was recently reconstituted in *A. thaliana* (12). In this line, a proto-starch sheath was produced by the introduction of both SAGA1 and SAGA2 (11). Starch granules accumulated around the matrices in a manner reminiscent of the *sbe1* mutant where starch granules fail to preferentially grow circumferentially to encapsulate the matrix and instead sit along the matrix as amorphously shaped granules. These observations indicate that additional factors are required for the proper circumferential growth of proto-pyrenoid associated starch granules in *A. thaliana*. Our results suggest that the expression of SBE1 could lead to circumferential growth of these proto-pyrenoid-associated starch granules, with the potential to enable the formation of a complete starch sheath around an engineered pyrenoid matrix.

Beyond pyrenoid assembly, our findings offer insights into the broader principles of how starch granules are directly shaped by proteins and may inform efforts to engineer starch morphology in other systems. Direct shaping of the lenticular starch granules in *A. thaliana* leaves has been demonstrated to occur via the N-terminal domain of STARCH SYNTHASE 4 (28, 37, 47) which is believed to localize the protein to distinct locations along the thylakoids (28, 48). In *Solanum tuberosum* amyloplasts, starch shape was found to be conferred by the starch-binding protein PROTEIN TARGETING TO STARCH2b (49). Our characterization of SBE1 broadens the range of possible mechanisms for direct starch granule shaping by proteins to include not only localized starch synthases and starch-binding proteins but also localized starch branching enzymes. Additionally, our findings suggest that starch granules can be directly shaped to match the contour of another cellular structure—in this case, the phase-separated pyrenoid matrix—by localizing a starch biosynthesis enzyme to that structure.

## Materials and Methods

### Chlamydomonas strains and culturing conditions

All strains generated and used in this study are listed in *SI Appendix*, Table S1. Chlamydomonas cells were maintained at room temperature (∼22°C) on 1.5% agar plates prepared with Tris-acetate-phosphate (TAP) medium under ambient light as described in Crans et al., 2026 (10). Cells were grown in liquid culture prior to experiments by inoculating Tris-phosphate (TP) minimal medium (same composition as TAP but without acetate) with the appropriate strain. Cultures were supplied with cool white LED light at ∼175 μmol photons⋅m^−2^⋅s^−1^ in an orbital shaker (Infors) set to 23°C and 120 rpm. The air was enriched to 3% (v/v) CO2 for ∼1-2 days until the cell concentration reached ∼0.5-1 x 10^6^ cells⋅mL^−1^. Cultures remained either in 3% CO2 conditions until harvest for time course experiments starting with cells acclimated to high CO2 or were moved to a different orbital shaker with the same settings except only atmospheric air (0.04% v/v levels of CO2) was provided for the specified time until harvest. Cells were harvested for experiments at a concentration of approximately 1-3 x 10^6^ cells⋅mL^−1^ (mid-log phase) measured using a Countess II Automated Cell Counter (Life Technologies).

### Confocal Microscopy

Live cells were imaged using µ-Slide 2 well chambered coverslips (Ibidi) for time course experiments and µ-Slide 8 well chambered coverslips for general localization with a 2-3 mm TP minimal medium 1.5% agarose pad on top of cells. The lid was removed from the chamber to allow for gas exchange and the agarose pad covered with sterile anti-evaporation oil (Ibidi) to prevent desiccation. For high-to-low CO2 time courses, samples were prepared and imaging started within ∼20 minutes after removal from the high CO2 growth chamber. To image cells, a Nikon A1R point scanning confocal microscope was used with a ×100 magnification/1.45 NA oil objective. Z-stacks were collected every 0.3 µm to capture whole cells. Light was constantly provided using the illuminator of the microscope (∼60 μmol photons⋅m^−2^⋅s^−1^). For microscopy experiments looking only at Venus fusion proteins, Venus fluorescence was collected using a 514 nm excitation laser and 522-555 nm emission filter and chlorophyll autofluorescence was collected with the same excitation laser but a 601-676 nm emission filter. For microscopy experiments looking only at mCherry fusion proteins, mCherry fluorescence was collected using a 561 nm excitation laser and 570-620 nm emission filter and chlorophyll autofluorescence was collected with the same excitation laser but a 663-738 nm emission filter. For the colocalization of Venus and mCherry, the fluorescence was collected sequentially as follows: Venus using a 488 nm excitation laser and 500-550 emission filter, mCherry using a 561 nm excitation laser and 570-620 nm emission filter, and chlorophyll autofluorescence using a 561 nm excitation laser and a 663-738 nm emission filter. For time course imaging, images were processed after collection using Nikon NIS elements software to optimize image quality with the Denoise.ai and 3D Deconvolution tools. Images collected for the localization of SBE1-Venus were not processed due to a lower signal-to-noise ratio. All images were processed with Fiji software (50) to display appropriate contrast.

### Diurnal Synchronization of Cells

Cells were synchronized as in Ral et al., 2006 (32) under photoautotrophic conditions (TP minimal medium) in low (0.04% v/v) CO2 by applying a 12-h-light/ 12-h-dark regimen for at least 7 days prior to harvest for microscopy. Cells were supplied with cool white LED light at ∼120 μmol photons⋅m^−2^⋅s^−1^ in flasks with stirring at 200 RPM. Cultures were never allowed to grow over 4 x 10^6^ cells⋅mL^−1^ and were diluted accordingly into fresh medium during the light period. Cultures destined for imaging were wrapped in aluminum foil 8 h into the last light period and harvested at a concentration of ∼1-2 x 10^6^ cells⋅mL^−1^ after the extended 16 h dark period. Samples were prepared for microscopy as described above in a dark room (<0.01 μmol photons⋅m^−2^⋅s^−1^) ∼30 minutes prior to the end of the dark period and were transported to the microscope in a dark box such that imaging would begin when the dark period would normally end. The samples were set up and brought into focus on the microscope under minimal light in ∼15-30 seconds before imaging.

### Cloning and Recombineering

Plasmids generated in this study are listed in *SI Appendix*, Table S3 and their sequences can be found in Datasets S2-4. pLM161-SBE1-Venus-3xFLAG was generated using a recombineering-based approach as previously described (51). The bacterial artificial chromosome containing the genomic sequence and ∼2.6kb of upstream sequences of the *SBE1* gene from Chlamydomonas as well as the acceptor plasmid and primers used to linearize the acceptor plasmid for recombination are listed in *SI Appendix*, Tables S2 and S3. The SBE1-Venus-3xFLAG construct was validated using whole plasmid Oxford Nanopore sequencing (Plasmidsaurus) which revealed six indel mutations in intron sequences and one in the upstream sequences preceding the *SBE1* gene when compared to the Chlamydomonas v.5.6 reference genome sequence. Three point mutations were also identified in intron sequences. All but one of these mutations were in long mononucleotide or dinucleotide repeats. We therefore deemed the construct acceptable for gene rescue and localization experiments.

To generate pMB-NATr-STA2-mCherry-6xHis, the *STA2* sequence from pLM005-Cre17.g721500-Venus-3xFLAG (4), the *mCherry-6xHis* and pLM006 backbone sequence from pLM006 (1), and the nourseothricin resistance gene (*NAT*) sequence from pAB262-NAR-R (52) were amplified by PCR (primers can be found in *SI Appendix*, Table S2) to generate 20 bp overhangs for NEBuilder® HiFi DNA Assembly. To generate pMB-STA2-mCherry-6xHis-F2A-NATr, the *STA2* sequence from pLM005-Cre17.g721500-Venus-3xFLAG (4), the *mCherry-6xHis* sequence from pLM006 (1), and the *NAT* sequence and plasmid backbone from pAB262-NAR-R (52) were amplified by PCR to generate 20 bp overhangs for NEBuilder® HiFi DNA Assembly with a gBlock (IDT) sequence coding for the F2A peptide (*SI Appendix*, Table S2). The fragments were assembled such that *NAT* was positioned after the *STA2-mCherry-6xHis* sequence and both sequences linked in-frame with the F2A peptide sequence between them. Constructs were validated using whole plasmid Oxford Nanopore sequencing (Plasmidsaurus).

### Chlamydomonas Transformations

Chlamydomonas transformations were performed as described in Hennacy et al., 2024 (6) with the following deviations. Cells were grown in TAP medium in an orbital shaker as described above in low (0.04% v/v) CO2 conditions to a concentration of ∼1-2 x 10^6^ cells⋅mL^−1^. 115 µL of the Chlamydomonas strain to be transformed and was combined with 5 µL of heat-denatured carrier DNA (MB grade from fish sperm, 10 mg/mL, Roche) and 5 µL of the linearized plasmid (see *SI Appendix*, Table S3 for the restriction enzyme used) to be transformed on ice for ∼5-10 minutes prior to electroporation. Transformants were selected on 1.5% agar TAP plates supplemented with either hygromycin or nourseothricin and were supplied with dim light (<10 μmol photons⋅m^−2^⋅s^−1^) for 3 days followed by ∼100 μmol photons⋅m^−2^⋅s^−1^ of cool white LED light for ∼7-10 days until colonies were of sufficient size. Successful transformants were screened by growing individual colonies in TP minimal medium for ∼1-2 days under low (0.04% v/v) CO2 conditions supplied with ∼100 μmol photons⋅m^−2^⋅s^−1^ of cool white LED light in 96-well plates followed by confocal microscopy to confirm the expression of the inserted construct.

### Transmission Electron Microscopy

For TEM, cells were cultured as described above under high (3% v/v) CO2 conditions and then transferred to grow in low (0.04% v/v) CO2 conditions for 4 hours prior to preparation. Cells were harvested at a concentration of ∼1-2 x 10^6^ cells⋅mL^−1^. Sample preparation was carried out as previously described (53). Briefly, cells were first fixed with glutaraldehyde, stained with OsO4, and finally embedded in Quetol epoxy resin. Thin (∼70 nm) sections were cut from the cured resin blocks and mounted on copper mesh grids as previously described (53). Samples were imaged using a Talos L120C G2 (S)TEM (Thermo Fisher Scientific) at the Imaging and Analysis Center, Princeton University.

### Analysis of Starch Sheath Granules in TEM Images

Analysis of the circumferential-to-radial length ratio of pyrenoid matrix adjacent starch granules in TEM images was carried out using the segmented line tool in Fiji software (50) to determine if there was a growth direction preference for starch sheath granules in each strain. TEM images of whole cells were used for measurement. For each starch granule in contact with the pyrenoid matrix, the length of the granule along the surface of the pyrenoid matrix was taken as the circumferential length and the perpendicular length of the granule from the midpoint of the circumferential line was taken as the radial length. The circumferential-to-radial length ratios were calculated by dividing the circumferential length by the radial length of a given granule. To test for significance between each strain, a Kruskal–Wallis test followed by Dunn’s multiple comparisons test with Bonferroni correction was performed on the ratios using R. Plots were generated for visualization using ggplot2 (geom_beeswarm) in R.

### Zymography

Cultures for zymography were inoculated into TP-minimal medium or TAP medium and grown as described above under high (3% v/v) CO2 conditions and then transferred to low (0.04% v/v) CO2 conditions for ∼21 hours. Cells were harvested as described in Courseaux et al., 2023 (38) with the following modifications. Cells were centrifuged at 600 x g for 5 minutes at 4°C, washed once in lysis buffer (50 mM HEPES pH 7.0, 10% glycerol, and cOmplete™, EDTA-free Protease Inhibitor Cocktail), centrifuged as before, and then resuspended in 1 mL lysis buffer per 50 mL original culture volume before being immediately flash frozen in liquid nitrogen for storage at -80°C. Lysate was prepared as follows. The frozen cells were thawed on ice and sonicated on ice for 2 minutes (cycles of 5 second pulse at 60% amplitude, 5 second rest) using a probe sonicator (QSonica). Lysis was confirmed using a microscope. The crude lysate was centrifuged at 10,000 x g for 20 minutes at 4°C. The resulting clarified lysate was transferred to a clean tube on ice and the protein concentration estimated using a NanoPhotometer® N50.

Native zymography for the detection of starch branching activity (44) based on phosphorylase A stimulation (43, 45) was carried out as described in Courseaux et al., 2023 (38) with the following changes. 50 µg of lysate from the above procedure was immediately loaded onto 7.5% TGX gels (Bio-Rad) in native sample buffer (Bio-Rad) and ran at 15 V·cm^-1^ for 3 h at 4°C in Tris-Glycine running buffer (Bio-Rad). Following electrophoresis, the gels were placed in 20 mL of lysis buffer without protease inhibitor with gentle shaking for 20 minutes. After washing, the gels were incubated for 16 hours at 4°C in 20 mL lysis buffer without protease inhibitor supplemented with 50 mM glucose-1-phosphate (Sigma), 2.5 mM AMP (Thermo Fisher Scientific), and 28 units of phosphorylase a from rabbit muscle (Sigma) with gentle shaking. Branching enzyme activity was revealed by staining with Lugol’s solution (0.1% I2, 1% KI) which stains highly branched α-glucans such as glycogen and amylopectin a reddish-brown (44).

### Immunoprecipitation Mass Spectroscopy

Cultures for immunoprecipitation and mass spectroscopy of wild-type cells expressing *SBE1-Venus-3xFLAG* were prepared as described previously (10). The protocol for the immunoprecipitation and mass spectroscopy was carried out as previously described (54). The Venus-3xFLAG data in Dataset S1 was previously published as a control in Hennacy et al., 2024 (6), which is licensed under CC BY 4.0.

### Disclosure of Delegation to Generative AI

The authors declare the use of generative AI (GAI) in the research and writing process. According to the GAIDeT taxonomy (2025), the following tasks were delegated to GAI tools under full human supervision: Proofreading and editing.

The GAI tools used were: Gemini 3.1 Pro, Gemini 1.5 Flash, and ChatGPT Pro 5.6. The GAI tools were used as follows: Gemini 3.1 Pro and 1.5 Flash were used to improve the phrasing of several sentences, and ChatGPT Pro 5.6 was used to double-check proper figure references and citations.

Responsibility for the final manuscript lies entirely with the authors. GAI tools are not listed as authors and do not bear responsibility for the final outcomes. Declaration submitted by: Micah I. Burton and Martin C. Jonikas.

## Acknowledgments

We thank Sophie Skanchy for writing code to assign CreIDs to the accession numbers in the unprocessed mass spectrometry data and feedback on the manuscript; Victoria L. Crans for manuscript feedback, advice and helpful discussions, Aastha Garde and all other present and former members of the Jonikas Lab, for advice and helpful discussions; Alistair J. McCormick, Luke C.M. Mackinder, and Lara Esch for helpful discussions and manuscript feedback; Shan He for manuscript feedback; Marie Bao, as part of Life Science Editors, for manuscript editing. Research reported in this publication was supported by the National Institute of General Medical Sciences (NIGMS) of the National Institutes of Health under grant numbers T32GM007388 and 1R01GM140032-01; by National Science Foundation Grant MCB-2410354; by the Bill and Melinda Gates Foundation and United Kingdom Foreign, Commonwealth & Development Office grant INV-054558; by the Howard Hughes Medical Institute; and by a grant to Princeton University from the Howard Hughes Medical Institute through the Gilliam Fellows Program. We acknowledge the use of Princeton’s Imaging and Analysis Center, which is partially supported by the Princeton Center for Complex Materials, a National Science Foundation Materials Research Science and Engineering Center (MRSEC; DMR-1420541). Confocal imaging was conducted with support from Gary Laevsky and Sha Wang at the Light Microscopy Facility, a Nikon Center of Excellence, in the Department of Molecular Biology at Princeton University. The content is solely the responsibility of the authors and does not necessarily represent the official views of the funders.

## Author Contributions

M.I.B. and M.C.J. designed the project; M.I.B. performed PCRs, transformations, confocal microscopy, cloning, and zymography; H.W. performed TEM preparation and imaging; L.W. performed immunoprecipitation-mass spectrometry; J.H. performed preliminary TEM. M.I.B. and M.C.J. analyzed the data; M.I.B. and M.C.J. wrote the paper; all authors contributed to editing the paper.

