## Supporting Information for "SBE1 drives the circumferential growth of the starch sheath around the *Chlamydomonas reinhardtii* pyrenoid"

**This PDF file includes:**

Figures S1 to S5
Tables S1 to S3
Legends for Movies S1 to S7
Legends for Datasets S1 to S4
SI References

**Other supporting materials for this manuscript include the following:**

Movies S1 to S7
Datasets S1 to S4

**Figures**

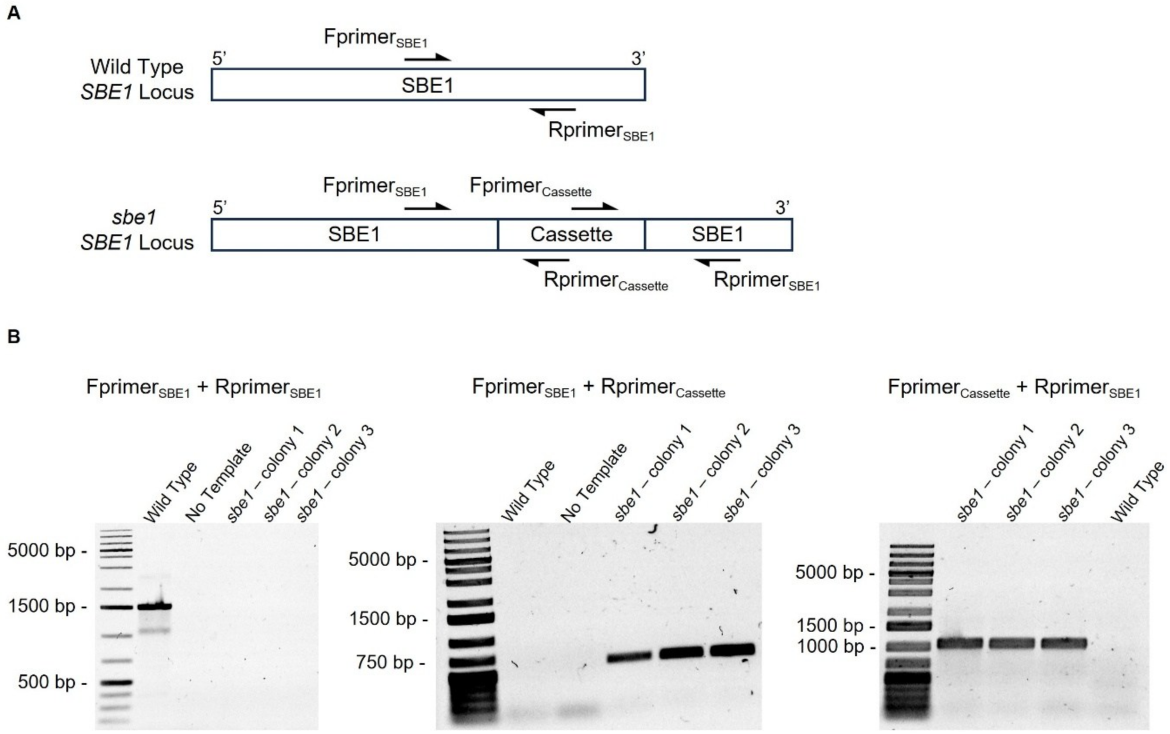

**Fig. S1.** The CLiP library s*be1* mutant LMJ.RY0402.151042 contains an insertional cassette in the *SBE1* gene. (A) Diagram of the PCR strategy used to confirm CLiP mutants. (B) The expected size for the Fprimer$_{SBE1}$/Rprimer$_{SBE1}$ primer pair **is** 1499 bp. The lack of a band in the *sbe1* lanes indicates that there is likely a disruption at the *SBE1* locus in the mutant. The expected sizes for the Fprimer$_{SBE1}$/Rprimer$_{Cassette}$ and Fprimer$_{Cassette}$/Rprimer$_{SBE1}$ primer pairs are 688 bp and 1068 bp respectively. The presence of the correctly-sized band in the *sbe1* lanes and its absence in the Wild Type lanes indicates the presence of the insertional cassette at the *SBE1* locus in the *sbe1* mutant.

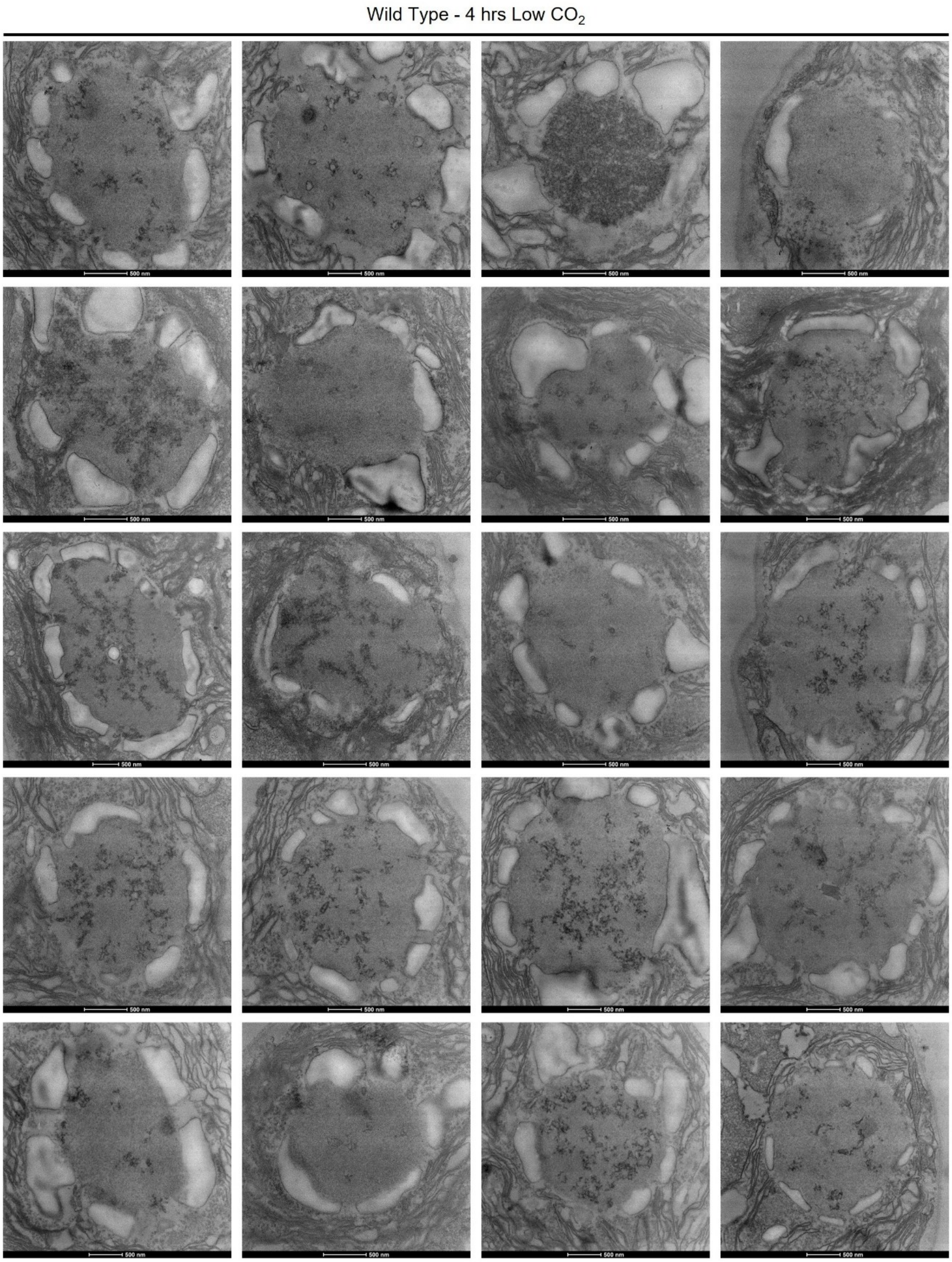

**Fig S2.** TEM images of pyrenoids from wild-type cells grown in TP minimal media in high (3% v/v) $CO_2$ levels and shifted to low (0.04% v/v) $CO_2$ levels for 4 hours before harvesting. All starch granules in contact with the pyrenoid matrix in the imaged pyrenoids were used in the analysis in Fig. 4*C.*

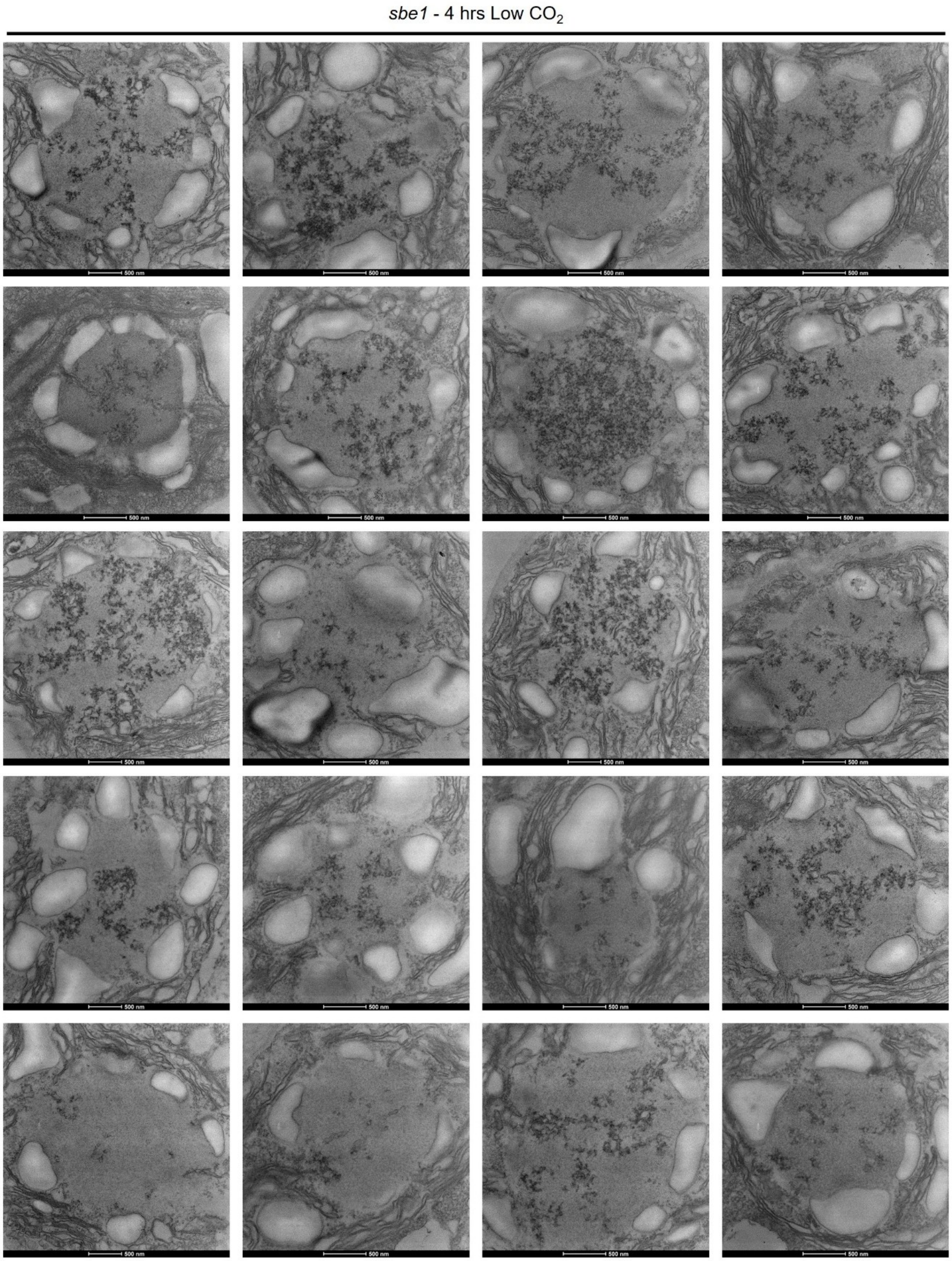

**Fig. S3.** TEM images of pyrenoids from *sbe1* mutant cells grown in TP minimal media in high (3% v/v) $CO_2$ levels and shifted to low (0.04% v/v) $CO_2$ levels for 4 hours before harvesting. All starch granules in contact with the pyrenoid matrix in the imaged pyrenoids were used in the analysis in Fig. 4*C.*

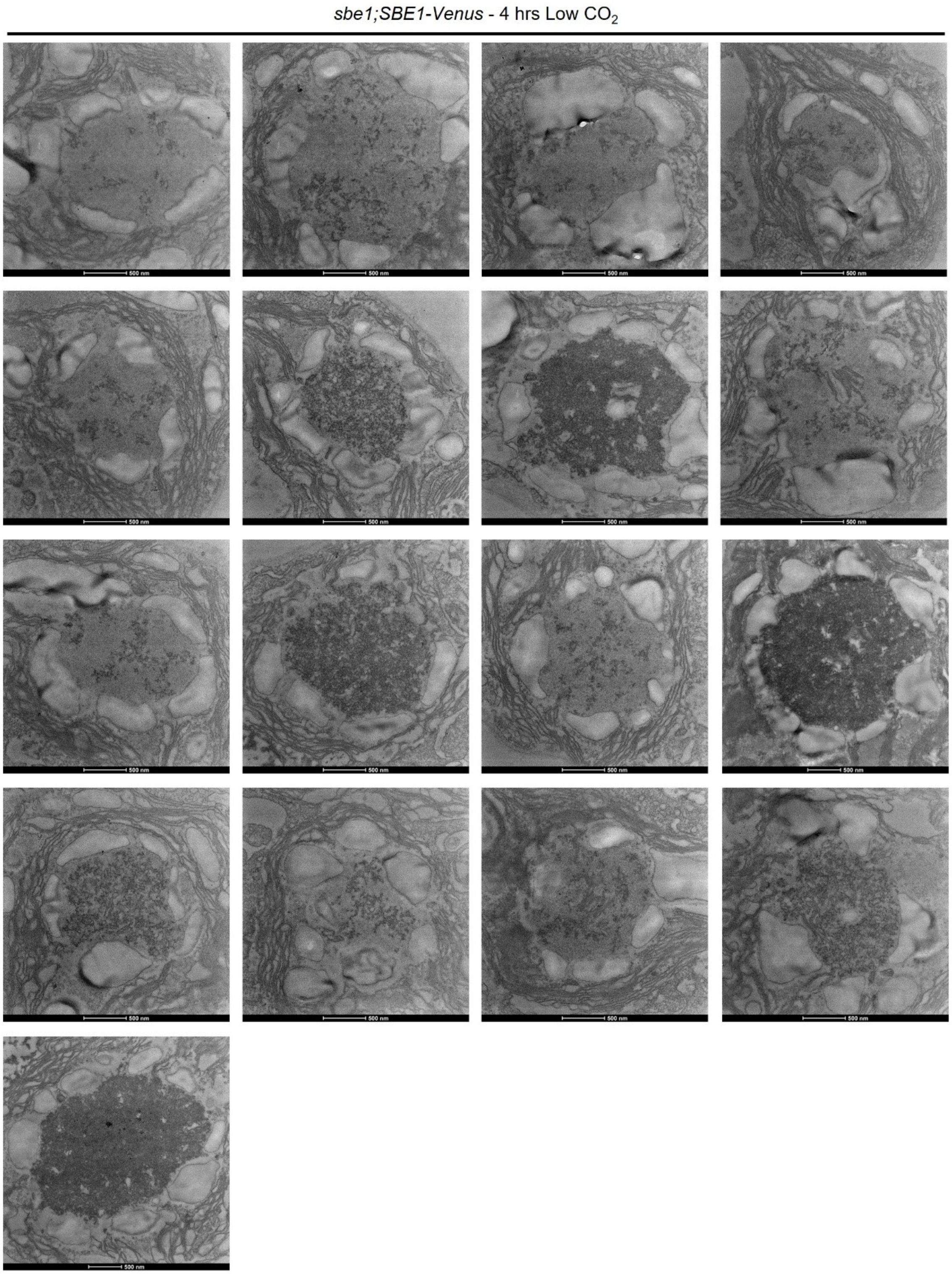

**Fig. S4.** TEM images of pyrenoids from *sbe1*;SBE1-Venus cells grown in TP minimal media in high (3% v/v) $CO_2$ levels and shifted to low (0.04% v/v) $CO_2$ levels for 4 hours before harvesting. All starch granules in contact with the pyrenoid matrix in the imaged pyrenoids were used in the analysis in Fig. 4*C.*

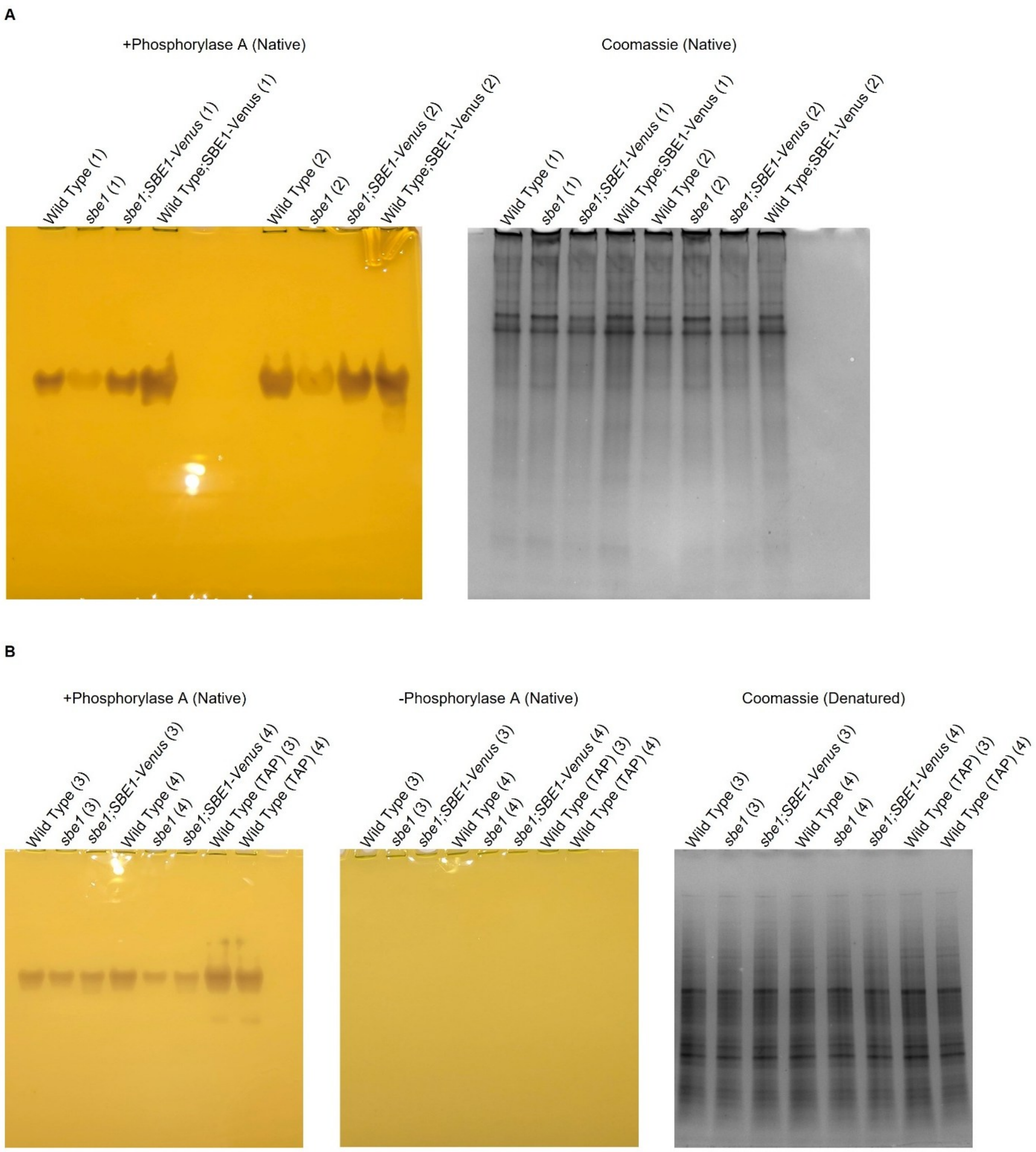

**Fig. S5.** Native zymograms using polyacrylamide gels containing no polysaccharide to indirectly assay starch branching activity in lysate. Cultures were grown at high (3% v/v) $CO_2$ and shifted to low (0.04% v/v) $CO_2$ for 21 hours prior to harvest. After running, zymogram gels were incubated for 16 hours at 4°C in assay buffer with gentle shaking with or without phosphorylase A prior to staining with Lugol's solution (1% KI, 0.1% $I_2$). Coomassie gels are shown to confirm that the measured

NanoPhotometer® concentrations used to calculate the amount of protein to load yielded similar amounts of protein between samples. Coomassie gels were stained immediately after running. (A) Full gel from Fig. 3*D*. Lysate was harvested from wild-type, *sbe1* mutant, *sbe1;SBE1-Venus*, and wild-type;SBE1-Venus cells grown in TP minimal media. (B) Lysates from cells grown in TP minimal media lacked several bands of branching activity that were previously reported for TAP-grown cells (1). Therefore, we harvested lysates from wild-type, *sbe1* mutant, *sbe1;SBE1-Venus* cells grown in TP minimal media, and TAP-grown wild-type cells to check if there were additional bands of activity in the TAP grown lysates compared to TP. 50 µg of protein was loaded in each labeled well except for the Coomassie stained gel in B where 12.5 µg was loaded in each well. Numbers in parentheses indicate replicate lysates from independent cultures.

**Table S1.** Chlamydomonas strains used in this study. (*) Strains that tended to clump in liquid culture. (**) Antibiotic used to select for positive transformants.

| Chlamydomonas Resource Center ID | Strain Description | Source | Antibiotic Resistance |
|---|---|---|---|
| CC-4533 | CMJ030 (Wild type) | Wild-type parent strain to the CLiP library (2) | None |
| CSI_FC1G03 | CMJ030;*STA2-Venus* | Wild-type strain expressing *STA2-Venus-3xFLAG* from the Chlamydomonas Spatial Interactome collection (3) | Paromomycin |
| LMJ.RY0402.151042 | *sbe1** | CLiP library (4) | Paromomycin |
| | *sbe1;SBE1-Venus** | *sbe1* mutant transformed with pLM161-SBE1-Venus-3xFLAG | Paromomycin, Hygromycin** |
| | CMJ030;*SBE1-Venus* | Wild-type strain CC-4533 transformed with pLM161-SBE1-Venus-3xFLAG | Hygromycin |
| | CMJ030;*STA2-mCherry* | Wild-type strain CC-4533 transformed with pMB-STA2-mCherry-6xHis-F2A-NATr | Nourseothricin |
| | *sbe1;STA2-mCherry** | *sbe1* mutant transformed with pMB-STA2-mCherry-6xHis-F2A-NATr | Paromomycin, Nourseothricin** |
| | *sbe1;SBE1-Venus;STA2-mCherry** | *sbe1;SBE1-Venus* transformed with pMB-NATr-STA2-mCherry-6xHis | Paromomycin, Hygromycin, Nourseothricin** |
| | *sta6;SBE1-Venus* | CC-5374 *sta6* starchless mutant transformed with pLM161-SBE1-Venus-3xFLAG | Hygromycin |

**Table S2.** PCR primers used in this study.

| Name | Use | Sequence | Source |
|---|---|---|---|
| Fprimer$_{SBE1}$ | Confirmation of CLIP mutant LMJ.RY0402.151042 | AAACAGAATGCGGTGTCCTC | Integrated DNA Technologies (IDT) |
| Rprimer$_{SBE1}$ | Confirmation of CLIP mutant LMJ.RY0402.151042 | TATAAAGCCGCACACAGCAC | IDT |
| Fprimer$_{Cassette}$ oMJ944 | Confirmation of CLIP mutant LMJ.RY0402.151042 | GACGTTACAGCACACCCTTG | Previously published (4) |
| Rprimer$_{Cassette}$ oMJ913 | Confirmation of CLIP mutant LMJ.RY0402.151042 | GCACCAATCATGTCAAGCCT | Previously published (4) |
| Fprimer$_{pLM161\text{-}SBE1}$ | Addition of homology arms for recombination of *SBE1* into pLM161 | CGACGCAGACCGCGGGCGGGCTTAGCGTCGGCAGCGGCGACGAAGTGGTGGGAGATCTGGGTGGCTCCG | Previously published (5) |
| Rprimer$_{pLM161\text{-}SBE1}$ | Addition of homology arms for recombination of *SBE1* into pLM161 | ATACGATGTCTATGTTCATGTCACGCAGCGAAAAGGAAGAGGTTACGGAGGAAGATCCTTTGATCTTTTCTACGGG | Previously published (5) |
| oMB143 (Forward) | Amplification of *STA2* out of pLM005-Cre17.g721500-Venus-3xFLAG | GGGACCTGATGGTGTTGGTG | IDT |
| oMB144 (Reverse) | Amplification of *STA2* out of pLM005-Cre17.g721500-Venus-3xFLAG | CCGGAGCCACCCAGATCTCC | IDT |
| oMB145 (Forward) | Amplification of *mCherry-6xHis* and pLM006 backbone out of pLM006 | GGAGATCTGGGTGGCTCCGG | IDT |
| oMB146 (Reverse) | Amplification of *mCherry-6xHis* and pLM006 backbone out of pLM006 | GTTTGCGGGTTGTGACTGAA | IDT |
| oMB147 (Forward) | Amplification of *crNAT* out of pAB262-NAR-R and the addition of homology arms for assembly with pLM006 backbone | TTCAGTCACAACCCGCAAACATGGGCACCACCCTGGACGA | IDT |
| oMB148 (Reverse) | Amplification of *crNAT* out of pAB262-NAR-R | CACCAACACCATCAGGTCCC | IDT |
| oMB149 (Forward) | Amplification of *STA2* out of pLM005-Cre17.g721500-Venus-3xFLAG | ACCAATCGTCACACGAGCCC | IDT |
| oMB150 (Reverse) | Amplification of *crNAT* and pAB262-NAR-R backbone out of | GGGCTCGTGTGACGATTGGTTTGTCTGCTCCCGGCATCCG | IDT |

| | | | |
|---|---|---|---|
| | pAB262-NAR-R and the addition of homology arms for assembly with STA2 from pLM005-Cre17.g721500-Venus-3xFLAG | | |
| oMB151 (Forward) | Amplification of *crNAT* and pAB262-NAR-R backbone out of pAB262-NAR-R and the addition of homology arms for assembly with gBlock mCherry-6xHis-F2A | TGGAGAGCAACCCGGGCCCC ATGGGCACCACCCTGGACGA | IDT |
| oMB152 (Reverse) | Amplification of *mCherry-6xHis* and pLM006 backbone out of pLM006 | ACAGCTCGTCCATGCCGCCG | IDT |
| gBlock mCherry-6xHis-F2A | Synthesized gBlock containing the partial 3' sequence from the *mCherry-6xHis* sequence in pLM006 and the F2A peptide sequence for the construction of pMB-STA2-mCherry-6xHis-F2A-NATr. | GCATGGACGAGCTGTACAGAT CTGGCGGTGGCCACCACCAT CACCACCACGGAGATCTCGT GAAGCAGACCCTGAACTTCGA CCTGCTGAAGCTGGCGGGCG ACGTGGAGAGCAACCCGGGC CCC | IDT |

**Table S3.** Plasmids used in this study. The listed restriction enzyme refers to the enzyme used to linearize the plasmid prior to electroporation into Chlamydomonas. For sequence maps of the plasmids generated in this study see Datasets S2-S4.

| Plasmid | Description/ Source | Antibiotic Resistance (Chlamydomonas) | Antibiotic Resistance (*E. coli*) | Restriction Enzyme |
|---|---|---|---|---|
| 12A3 | BAC containing the SBE1 genomic sequence (Chlamydomonas Resource Center) | N/a | Chloramphenicol | N/a |
| pLM161 | Recombineering backbone plasmid (5) | Hygromycin | Kanamycin | N/a |
| pLM161-SBE1-Venus-3xFLAG | SBE1 genomic sequence from BAC 12A3 with an additional 2613 bp upstream of the start codon recombineered into pLM161 | Hygromycin | Kanamycin | I-SceI |
| pLM005-Cre17.g721500-Venus-3xFLAG | Previously published (3). Contains *STA2-Venus-3xFLAG*. | Paromomycin | Ampicillin | N/a |
| pLM006 | Previously published (6). Contains *mCherry-6xHis*. | Hygromycin | Ampicillin | N/a |
| pAB262-NAR-R | Previously published (7). Contains *NAT*. | Nourseothricin | Ampicillin | N/a |
| pMB-NATr-STA2-mCherry-6xHis | NEBuilder® HiFi DNA Assembly of PCR amplified *STA2* from pLM005-Cre17.g721500-Venus-3xFLAG, *mCherry-6xHis* from pLM006, and *NAT* from pAB262-NAR-R. Backbone from pLM006. | Nourseothricin | Ampicillin | EcoRV |
| pMB-STA2-mCherry-6xHis-F2A-NATr | NEBuilder® HiFi DNA Assembly of PCR amplified *STA2* from pLM005-Cre17.g721500-Venus-3xFLAG, *mCherry-6xHis* from pLM006, *NAT* from pAB262-NAR-R, and a gBlock containing the F2A peptide. | Nourseothricin | Ampicillin | ZraI |

| | |
|---|---|
| | Backbone from pAB262-NAR-R. |

**Movie S1 (separate file).** Movie of the cells in Fig. 1*B*. The cells were grown in TP minimal media at high (3% v/v) $CO_2$ then transferred to a microscopy slide at air (0.04% v/v) levels of $CO_2$ prior to the first acquisition. Cells were imaged on TP-agarose pads under continuous illumination. Images were acquired every 15 minutes. Green signal is from STA2-Venus. Magenta signal is from chlorophyll autofluorescence.

**Movie S2 (separate file).** Movie of the dividing cell from Fig. 2 dividing together with another cell. Time course of cell division of a cell expressing STA2-Venus grown in TP minimal media at high (3% v/v) $CO_2$ then transferred to a microscopy slide at air (0.04% v/v) levels of $CO_2$ for 5 hours prior to the first acquisition. Images were acquired every 15 minutes. Time point 0 min indicates when full cleavage of the chlorophyll autofluorescence occurs. Green signal is from STA2-Venus. Magenta signal is from chlorophyll autofluorescence. Cells were imaged on TP-agarose pads and supplied with continuous illumination.

**Movie S3 (separate file).** Movie of a dividing cell where one daughter inherits nearly all starch sheath granules. Time course of cell division of a cell expressing STA2-Venus grown in TP minimal media at high (3% v/v) $CO_2$ then transferred to a microscopy slide at air (0.04% v/v) levels of $CO_2$. Images were acquired every 10 minutes. Time point 0 min indicates when full cleavage of the chlorophyll autofluorescence occurs. Green signal is from STA2-Venus. Magenta signal is from chlorophyll autofluorescence. Cells were imaged on TP-agarose pads and supplied with constant illumination.

**Movie S4 (separate file).** Movie of the diurnally entrained cell from Fig. 3. Time course of starch plate growth in cells expressing STA2-Venus grown in TP minimal media at air (0.04% v/v) levels of $CO_2$ . Cells were grown under a 12h-light/ 12h-dark regiment for at least 7 days before being subjected to a shortened day of 8 hours of light followed by 16 hours of darkness. Cells were imaged on TP-agarose pads and supplied with constant illumination. Images were acquired every 15 minutes. Green signal is from STA2-Venus.

**Movie S5 (separate file).** Expanded view of *sbe1* mutant cells from the experiment in Fig. 4*E-I*. Cells were grown in TP minimal media at high (3% v/v) $CO_2$ then transferred to a microscopy slide at air (0.04% v/v) levels of $CO_2$. Images were acquired every 20 minutes. The first part of the movie is a scroll through the z-stack at time = 0 min. The time course then plays showing the max-Z projections of each time point. The movie ends with a second scroll through the z-stack at time = 240 min. Green signal is from STA2-mCherry. Magenta signal is from chlorophyll autofluorescence. Cells were imaged on TP agarose pads and supplied with continuous illumination.

**Movie S6 (separate file).** Expanded view of *sbe1;SBE1-Venus* cells from the experiment in Fig. 4*J-N*. Cells were prepared as in Movie S5. The file plays as in Movie S5.

**Movie S7 (separate file).** Expanded view of wild-type cells from the experiment in Fig. 4*O-S*. Cells were prepared as in Movie S5. The file plays as in Movie S5.

**Dataset S1 (separate file).** Total spectral counts from a FLAG co-immunoprecipitation followed by mass spectrometry of cells expressing either SBE1-Venus-3xFLAG or Venus-3xFLAG. The Venus-3xFLAG data were previously published as a control in Hennacy et al., 2024 (8), which is licensed under CC BY 4.0. The SBE1-Venus-3xFLAG data were collected using the same protocol. Spectral counts are color coded along a spectrum where red indicates high counts and white indicates low counts. The alternate IDs of the five proteins listed in Fig. 5*D* are highlighted in yellow for ease of reference.

**Dataset S2 (separate file).** Sequence map for pLM161-SBE1-Venus-3xFLAG.

**Dataset S3 (separate file).** Sequence map for pMB-NATr-STA2-mCherry-6xHis.

**Dataset S4 (separate file).** Sequence map for pMB-STA2-mCherry-6xHis-F2A-NATr.
